# An Evaluation of the Efficacy of Single-Echo and Multi-Echo fMRI Denoising Strategies

**DOI:** 10.1101/2025.08.28.672985

**Authors:** Toby Constable, Jeggan Tiego, Kane Pavlovich, Arshiya Sangchooli, Priscilla Thalenberg Levi, Bree Hartshorn, Jessica Kwee, Kate Fortune, Kate Cooper, Sam Brown, James McLauchlan, Nancy Ong Tran, Rebecca O’Neill, Mark A. Bellgrove, Alex Fornito

## Abstract

Resting-state functional magnetic resonance imaging (rsfMRI) is commonly used to study brain-wide patterns of inter-regional functional coupling (FC). However, the resulting signals are vulnerable to multiple sources of noise, such as those related to non-neuronal physiological fluctuations and head motion, which can alter FC estimates and influence their associations with behavioral outcomes. The best strategy for acquiring and processing rsfMRI data to mitigate noise remains an open question. In this study of 358 healthy individuals, we compared the denoising efficacy of 60 multi-echo (ME) and 30 single-echo (SE) rsfMRI preprocessing pipelines across six distinct measures of data quality. We also evaluated how each pipeline influences the effect sizes of FC-based predictive models of personality and cognitive measures estimated via cross-validated kernel ridge regression. We found that ME pipelines generally showed superior denoising efficacy to SE pipelines, but that no single pipeline was associated with both superior denoising efficacy and behavioural prediction. Using a heuristic scheme to rank pipelines across benchmark evaluations, we found that an ME acquisition combined with Automatic Removal of Motion Artifacts Independent Component Analysis (ICA-AROMA) and Regressor Interpolation at Progressive Time Delays (RIPTiDe) offered a reasonable compromise between denoising efficacy and brain-behavior predictions for both ME and SE data. In general, ME pipelines ranked more highly than SE pipelines. These findings support the use of ME acquisitions in future work but suggest that no single denoising pipeline should be considered optimal for all purposes.

## An Evaluation of the Efficacy of Single-Echo and Multi-Echo fMRI Denoising Strategies

Functional magnetic resonance imaging (fMRI) detects changes in blood (de)oxygenation to infer neuronal activity (Hillman, 2014; Ogawa et al., 1990). The resulting blood oxygenation level-dependent (BOLD) signal is heavily influenced by non-neuronal factors such as head motion and physiological noise, which can compromise measurement fidelity (Hillman, 2014; Liu, 2016; Ogawa et al., 1990) and thus complicate attempts to obtain reliable effect sizes for brain-behaviour associations (Marek et al., 2022). This concern is particularly acute in the analysis of resting-state inter-regional functional coupling (FC) estimates, which are notoriously susceptible to various sources of noise (Birn et al., 2014). Developing preprocessing strategies for more effective separation of signal from noise is essential for generating valid and replicable insights from these analyses.

Head-motion constitutes one of the most influential sources of noise in BOLD data (Aquino et al., 2020; Ciric et al., 2017; Parkes et al., 2018; Power et al., 2012; Satterthwaite et al., 2013; Van Dijk et al., 2012). A simple strategy for addressing motion-related confounds in fMRI analyses involves regressing the 6 standard head motion parameters describing rotations and translations measured during image realignment, their quadratic expansion, first-order derivatives, and the quadratics of the derivatives from the BOLD signal using a procedure dubbed 24P regression (Satterthwaite et al., 2013). To further account for signal fluctuations of a non-neuronal origin, it is also common to remove signals averaged from the cerebrospinal fluid (CSF) and white matter (WM) tissue compartments (which we term 2P regression), optionally also in combination with their expansion and derivative terms (which we collectively term 8P regression) (Aquino et al., 2020; Parkes et al., 2018). A common additional step involves performing global signal regression (GSR) or grey matter signal regression (GMSR), which involves removing the average signal taken from the entire brain or grey matter tissue compartment and, optionally, the corresponding expansion and derivative terms. Finally, some analyses involve censoring high-motion frames, as estimated using a measure of framewise displacement (FD) (Ciric et al., 2017; Parkes et al., 2018; Power et al., 2012), either by removing those frames or using delta function regressors (Satterthwaite et al., 2013). In some cases, an interpolation is applied to replace the missing data (Power et al., 2013; Wilke & Baldeweg, 2019).

The GSR and GMSR procedures have attracted controversy. They both centre the distribution of FC estimates on zero (Fox et al., 2009), giving rise to spurious anticorrelations (Murphy et al., 2009) and distorting group differences in FC estimates (Saad et al., 2012). The technique may also be overly aggressive, potentially removing signals of interest from the data (Chang & Glover, 2009). However, global signal fluctuations are highly correlated with non-neuronal physiological fluctuations (Power et al., 2017; Xifra-Porxas et al., 2024) and their removal improves the anatomical specificity of inter-regional FC patterns (Fox et al., 2009), strongly mitigates motion-related signal contamination (Ciric et al., 2017; Parkes et al., 2018), improves correlations between FC and behavioural measures (J. Li et al., 2019), and yields BOLD signals that correlate more strongly with underlying neuronal calcium dynamics (Matsui et al., 2016).

Alternative methods for addressing global signal artifacts (also called widespread signal deflections, or WSDs) that may address the documented limitations of GSR have been proposed. One approach, termed Diffuse Cluster Estimation and Removal (DiCER), uses an iterative clustering procedure to remove WSDs, demonstrating variable performance relative to GSR depending on the characteristics of the data and the specific parameters used in its implementation (Aquino et al., 2020; Pavlovich et al., 2024). Another approach uses temporal independent component analysis (ICA) to separate components that represent distinct WSDs (Glasser et al., 2019). Although this approach offers an elegant solution, it requires large amounts of data and is difficult to apply to many extant fMRI datasets. One method gaining popularity involves estimating spatially and temporally heterogeneous low-frequency oscillations (LFO) that are unlikely to have a neural origin directly from BOLD time-series data using spatially dependent time-lagged regression models, as implemented in the Regressor Interpolation at Progressive Time Delays (RIPTiDE) algorithm (Frederick et al., 2012). LFOs predominantly occur on the haemodynamic rather than neural time scale and propagate through the brain’s vasculature, suggesting that they may originate from changes in arterial pressure (Tong et al., 2019). Recent work has found that RIPTiDE can mitigate spurious inflations of FC that occur as a function of time spent in the scanner (functional connectivity inflation; FCI) (Korponay et al., 2024). GSR/GMSR can also reduce FCI but an advantage of the RIPTiDe algorithm is that it does not yield spurious anti-correlations by construction (Erdoğan et al., 2016) .

Another popular class of denoising strategies relies on spatial independent component analysis (s-ICA), which is a dimensionality reduction technique that decomposes spatiotemporal fMRI signals into multiple sources (independent components; ICs) with maximal spatial independence (McKeown et al., 2003). Each IC may represent noise, neuronal signal, or a mixture of both. These components can be classified and used to selectively remove contributions from spatially structured noise signals. For instance, ICA-based Automatic Removal of Motion Artifacts (ICA-AROMA) is a widely used automatic denoising method that classifies components as either signal or noise based on several data-driven heuristics, under the assumption that true signal components typically exhibit low-frequency power, regular oscillations, and predominantly localize to grey matter (Griffanti et al., 2017; Pruim et al., 2015). ICA-AROMA thus identifies noise components based on high-frequency content, correlation with head motion realignment parameters, and activity near the edges of the brain or within the cerebrospinal fluid (CSF) (Pruim et al., 2015).

A more elaborate ICA-based denoising technique is FMRIB’s Independent Component Analysis-based X-noiseifier (ICA-FIX) (Salimi-Khorshidi et al., 2014). This method first involves hand-classifying ICs as either “signal” or “noise” in a training set derived from a larger data pool. These labels then serve as a “ground truth” to train a classifier for automatically rejecting noise components for the rest of the dataset. This approach allows for more tailored classifications given the particulars of the data by leveraging human expertise to improve the denoising process (Griffanti et al., 2017). Signal components can be manually classified according to similar heuristics used by ICA-AROMA (e.g., high-frequency power, irregular oscillatory patterns, poor localisation to grey matter), but can also include more complex spatial and/or temporal properties that ICA-AROMA may miss (Griffanti et al., 2017).

A third ICA-based approach requires a multi-echo (ME) acquisition. Unlike single-echo (SE) acquisitions, ME fMRI captures multiple images per radiofrequency (RF) pulse, each with varying contrast levels (Kundu et al., 2012, 2017; Lynch et al., 2021; Posse et al., 1999). Early echoes exhibit higher signal intensity and lower drop-out due to their temporal proximity to the RF pulse, while later echoes provide greater tissue contrast but lower signal intensity, except for tissues with slower decay rates (Kundu et al., 2017). Applying a weighted average across each echo yields an optimally combined time-series in which signal-quality and tissue-contrast ratios are maximized and signal dropout is minimized. BOLD and non-BOLD signals decay in intensity after each RF pulse at a known T₂* rate, which can be compared to empirical T₂* measures. When combined with spatial independent component analysis (sICA), multi-echo (ME) acquisition can aid in distinguishing signal from noise based on the T₂* of each independent component (IC). Comparing the relative fits of known BOLD and non-BOLD T₂* trends to the empirical data yields two parameters: κ and ρ. The parameter κ represents the likelihood that a component contains BOLD-related information whereas ρ represents the likelihood of non-BOLD (noise) contributions. The ratio between these parameters assists in determining whether a component should be retained or rejected (Kundu et al., 2012). Both the optimal combination of the ME time series and the ME-ICA approach individually contribute to an increase in signal-to-noise ratio (SNR) and can be effectively integrated into the same preprocessing pipeline (Steel et al., 2022). Accordingly, some work indicates that ME protocols can yield similar data quality to long SE acquisitions with approximately half the scan length (Lynch et al., 2020). ME-based pipelines can successfully mitigate motion-related contamination but, on their own, do not completely remove the WSDs that are targeted by GSR/GMSR and related techniques (Power et al., 2018).

Several prior studies have compared the relative strengths and weaknesses of some of the common fMRI denoising strategies discussed above. For example, Ciric et al. (2017) compared 14 denoising pipelines for SE data across six metrics, observing considerable between-pipeline variability across various estimates of data quality. The authors concluded that GSR and volume censoring effectively mitigate noise resulting from head motion. Parkes et al. (2018) compared 19 pipelines for SE data. They also found considerable variability in denoising efficacy across pipelines and concluded that pipelines incorporating ICA-AROMA and GSR perform well at mitigating motion-related noise. Few direct comparisons of ME and SE acquisitions have been reported. In one analysis, Di Pasquale et al. (2017) compared several popular SE and ME data preprocessing strategies, and found that ME-ICA outperformed ICA-AROMA and ICA-FIX according to several established fMRI data quality. RIPTiDe has recently been found to remove sources of noise that more commonly used methods may miss (Korponay et al., 2024).

Most of these studies have used different measures of denoising efficacy or data quality that aim to quantify the level of residual noise (e.g., motion contamination) found in the data after denoising. Although useful, such measures only consider removal of noise and do not quantify preservation of signal of interest. This is important, since a given denoising method may be overly aggressive, removing all noise-related contamination along with neuronal variance. Several alternative approaches have been developed to investigate this question (Aquino et al., 2020; Glasser et al., 2019), but perhaps the simplest strategy involves an evaluation of the degree to which the processing pipeline influences the performance of an FC-based predictive model of some behavioural outcome. Using this approach, Li et al. (2019) showed that SE pipelines incorporating GSR generally yielded models that more accurately predicted distinct aspects of personality and cognition than pipelines omitting this step. Another recent study comparing a wider range of SE pipelines found that no single pipeline consistently excelled at denoising efficacy and behavioural prediction, but that combining ICA-FIX and GSR offered a reasonable balance between these two priorities (Pavlovich et al., 2024).

No study to date has compared different SE and ME data preprocessing strategies across both data quality measures and the ability to predict individual differences in behaviour. To this end, we compared 90 different denoising pipelines for single and multi-echo fMRI data, focusing on denoising efficacy and behavioural predictions with respect to seven different measures of personality and cognition. This approach allowed us to investigate whether there is a specific acquisition and processing pipeline that is simultaneously associated with optimal denoising efficacy and behavioural prediction performance.

## Method

### Participants

A total of 418 community-dwelling adults aged 18-45 were recruited for a broader project examining brain-behaviour relationships. Inclusion in this analysis was contingent on participants possessing fMRI and T1-weighted images. Participants were recruited via an online campaign targeting various social media and other outlets, coordinated by the private company, Trialfacts (https://trialfacts.com/). Written informed consent was required before study participation. Inclusion criteria for this study included aged 18-45 years; right-handedness; European ancestry (defined as all four grandparents of European descent); no history of frequent headaches or migraines, past seizures, concussions, or loss of consciousness that lasted more than 3 minutes; English as the first spoken language; normal or corrected-to-normal vision; no metal in the body; no history of neurosurgery; not currently pregnant or attempting to conceive; no history of receiving hormone blockers or hormone replacement; no history of gender-affirming surgery; no history of steroid abuse; no neurological illness; no history of significant head injury inducing loss of consciousness; and no previous experience of electroconvulsive therapy. Experience of psychiatric symptoms or past/current psychiatric diagnosis and/or treatment were also used as a basis for exclusion for the present analysis to minimise sample heterogeneity. Treatment in this context was operationalized as no transcranial magnetic stimulation (TMS) within the past 6 months; no more than 2 weeks of TMS exposure in the last year; no more than 3 months continuous TMS at any point of life; and no history of psychiatric diagnosis, hospitalization, or medication use. All participants provided informed consent in accordance with Monash Human Research Ethics Committee guidelines.

### Data Acquisition

ME-rs-fMRI data were collected on a Siemens 3T Skyra (Siemens Healthineers, Erlangen, Germany) using an interleaved sequence with the following parameters: isotropic voxel size = 3.2mm, slices = 40, repetition time (TR) = 0.91s, phase encoding direction = R-L, echo times TE1 = 12.60 ms, TE2 = 29.23 ms, TE3 = 45.86 ms, TE4 = 62.49 ms, flip angle = 56°, bandwidth = 2520 Hz/Px, echo spacing =0.00025, GRAPPA acceleration factor *R*= 2, Volumes = 767, Multi-band Factor = 4. An opposite phase encoding scheme (L-R) with 10 volumes, but otherwise identical parameters, was also acquired for susceptibility distortion correction. T1-weighted images were also acquired using a magnetization-prepared 2 rapid-acquisition gradient-echoes (MP2RAGE) MRI sequence (TR = 5000ms, TE = 2.98ms, TI1/T12 = 700/2500ms, 1.0mm cubic voxel size).

### Data Processing

Figure 1 depicts the analysis workflow for our comparative evaluation. The second echo image (i.e., TE = 29.23ms) was used for all SE preprocessing pipelines. The following provides details of the specific steps applied to the anatomical and functional data.

**Figure 1.**
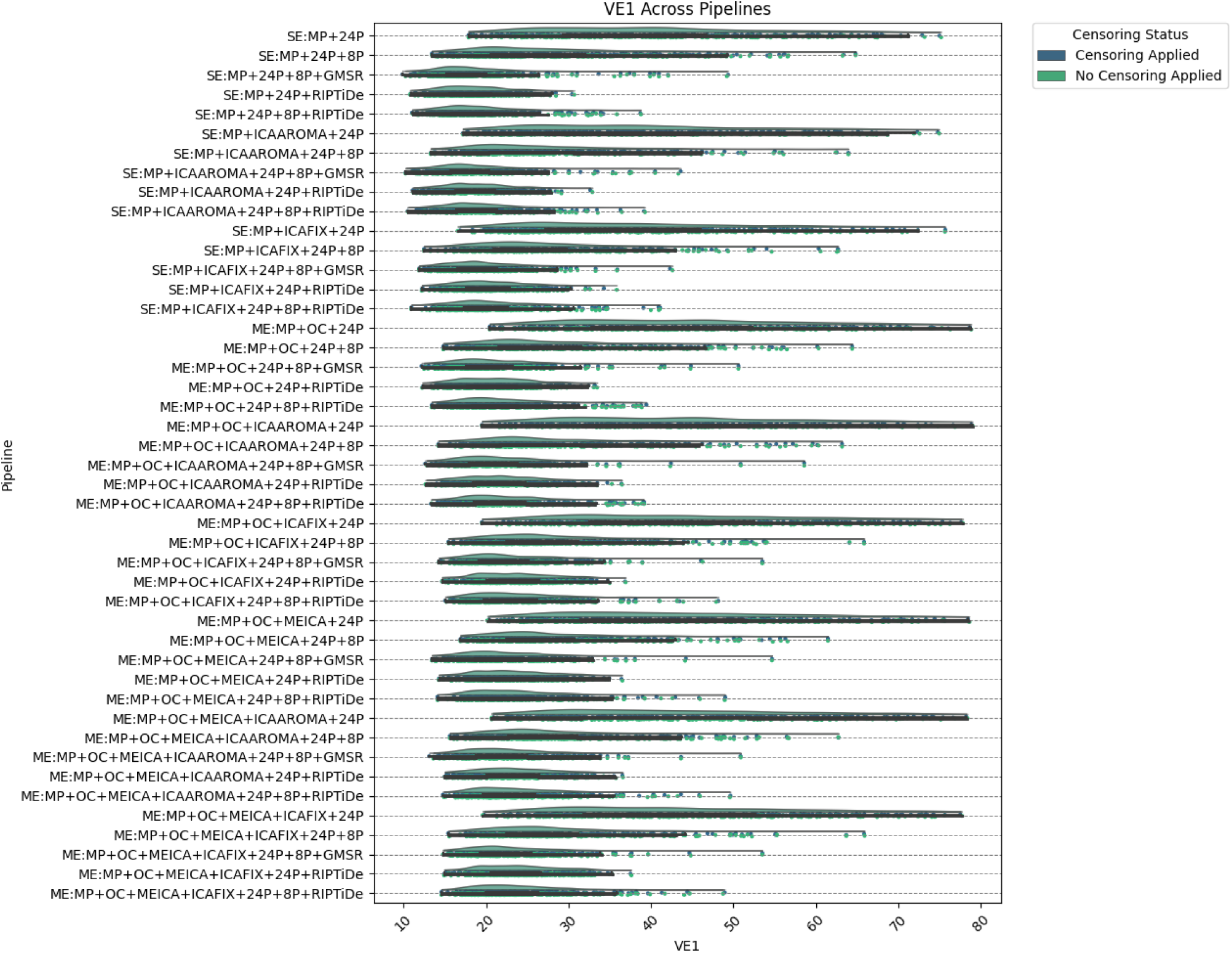
The effect of various denoising strategies on subject-level VE1, aggregated at the pipeline level. Smaller values indicate less noise. Note that histograms for Censoring Status (applied or not applied) overlap substantially. *MP* refers to minimal preprocessing with fMRIPrep.

### T1-weighted MP2Rage Processing

The MP2RAGE sequence involves the acquisition of two structural images with different inversion times (INV1 and INV2) which are then combined into a single T1-weighted image (UNI). The combination provides enhanced grey-to-white matter contrast and improves tissue segmentation accuracy (Marques et al., 2010; O’Brien et al., 2014) but can increase noise in non-brain regions, air, and large sinus spaces. To suppress such noise, we generated whole-brain masks and masks of several non-brain regions from the full INV2 image using the presurfer package (Kashyap et al., 2021) (https://github.com/srikash/presurfer). These masks were then used to strip regions affected by noise in the spatially aligned UNI image. Visual quality control confirmed the success of this approach. The processed full-head UNI image was then used as input to the Freesurfer recon-all pipeline for denoising, cortical reconstruction, and segmentation (Dale et al., 1999) (https://surfer.nmr.mgh.harvard.edu/). The resulting images were then incorporated into the fMRI processing pipelines, as detailed below.

### Functional Data Processing – minimal preprocessing (MP)

Functional MRI images were minimally preprocessed (MP) using the container fMRIPrep 23.1.3 (Esteban et al., 2019), based on Nipype 1.8.6 (Gorgolewski et al., 2011) (https://fmriprep.org/en/stable/). In brief, for anatomical data, this involved the creation of spatial maps quantifying the voxelwise probability of gray matter, white matter, or cerebrospinal fluid (Zhang et al., 2001), followed by registration to standard space (MNI152NLin2009cAsym) (Avants et al., 2008). Freesurfer masks and surfaces were used to refine brain-masks derived via antsBrainExtraction (Tustison et al., 2021) per fMRIPrep-specific sequences (i.e., from aseg.mgz). fMRI data were corrected for head motion via rigid body alignment (Jenkinson et al., 2002) and for susceptibility distortions using the reverse polarity image (Andersson et al., 2003; Smith et al., 2004). The functional scans were then co-registered to T1w space using boundary-based registration (Greve & Fischl, 2009) and spatially normalized to MNI152NLin2009cAsym space using non-linear registration (Avants et al., 2008). These same steps were applied independently to the fMRI time series obtained at each echo time, but parameters required for head motion correction and susceptibility distortion correction were estimated from the first echo and then applied to the remaining echo data, as per the default procedure in fMRIPrep. Slice timing correction was not performed due to the short TR of the fMRI acquisition sequence.

### ME Optimum Combination (OC)

A T2* map was estimated from the individually preprocessed echoes by fitting the maximal number of echoes with reliable signal for a given voxel to a monoexponential signal decay model using nonlinear regression and the calculated T2* map was then used to optimally combine preprocessed BOLD data across echoes, following Posse et al. (1999). Specifically, data across echoes were combined using a weighted average that prioritizes information from echo times near 30 ms for superior signal dropout to tissue contrast ratios. This step was executed using the software package *tedana* (DuPre et al., 2021) (https://tedana.readthedocs.io/en/stable/).

#### FSL-MELODIC

For SE pipelines, spatial ICA was run using the FSL Multivariate Exploratory Linear Optimized Decomposition into Independent Components (MELODIC) tool, performed on a 3mm spatially smoothed preprocessed second echo image (https://web.mit.edu/fsl_v5.0.10/fsl/doc/wiki/MELODIC.html).

For ME pipelines, the preprocessed, optimally combined BOLD images were smoothed using a 3mm Gaussian kernel and then subjected to spatial ICA with FSL-MELODIC (Beckmann & Smith, 2004), yielding a mixing matrix in which each column represents an estimated IC and rows represent volumes. Using this mixing matrix as input for all subsequent ICA-based denoising strategies ensured that all algorithms were parsing signal and noise using the same basis.

#### ICA-AROMA

The corresponding FSL-MELODIC outputs for SE and ME pipelines were used as input for ICA-AROMA (Pruim et al., 2015) (https://github.com/maartenmennes/ICA-AROMA). The algorithm was run using the reference BOLD image (a static image created via the median of motion-corrected volumes) created by fMRIPrep. The preprocessed T1w image created by fMRIPrep was skull-extracted using the FSL-Brain Extraction Tool (https://web.mit.edu/fsl_v5.0.10/fsl/doc/wiki/BET(2f)UserGuide.html). The resulting brain mask and skull-extracted images were also supplied to MELODIC. All images were spatially aligned using boundary-based registration (Greve & Fischl, 2009).

#### ME-ICA

The non-smoothed individually preprocessed echoes were processed with *tedana* for optimal combination and ME-ICA. ME-ICA applies automatic rejection of components based on the associated 𝜅/𝜌 ratio. These terms reflect the degree of model fit for known BOLD and non-BOLD T₂* trends with respect to the empirical data (Kundu et al., 2012). The MELODIC-estimated mixing matrix was used as input for the algorithm to ensure equivalence between pipelines. No smoothing was applied, as is standard for ME-ICA (Kundu et al., 2012).

#### ICA-FIX

ICA-FIX involves hand-classification of ICs estimated by FSL-MELODIC as either “signal” or “noise” within a training set derived from a larger collection of data. Hand-labelling of noise components was conducted twice for the same 20 subjects (10 male, 10 female) after FSL-MELODIC, once for their smoothed SE data and once for their smoothed optimally combined ME data. As such, separate training sets were generated for the SE and ME processing streams.

Manual classification was performed according to robust heuristics previously explicated in the literature (Griffanti et al., 2017). Leave-one-out cross-validation was used to assess classification accuracy, which involved iteratively training the classifier on all but one participant and using the left-out participant to evaluate the classifier’s alignment with manual classifications. This alignment was assessed over a variety of rejection thresholds, which determine how aggressive the algorithm is in identifying noise components. Subject-wise true positive rates (TPR) and true negative rates (TNR) were averaged across the 20 participants to estimate overall classifier performance for each rejection threshold. Previous literature suggests that TPR should approximate 90%, while TNR can be as low as 10%, to maintain the maximum amount of signal (Carone et al., 2017). For both our SE and ME datasets, a rejection threshold of 40 was identified as optimal for balancing TPR and TNR (SE: TPR [Mean, Median] = [87.70%, 88.10%]; TNR [Mean, Median] = [86.40%, 87.80%]; ME: TPR [Mean, Median] = [91.10%, 91.40%]; TNR [Mean, Median] = [89.00%, 89.90%]).

### Removing ICA-derived noise signals

For ME pipelines combining ICA-based denoising strategies (e.g. ME-ICA + ICA-AROMA), the list of noise components extracted via each denoising algorithm was combined into a single vector describing the noise component index positions (duplicates removed). The mixing matrix and the full list of rejected components was then supplied to FSL’s fsl_regfilt function for denoising via non-aggressive ordinary least squares regression.

### 8P, GMSR, and Spike Regression

Classical 8P, 8P+GMSR, and spike regression (SR), were applied in a second step, after ICA-based denoising, according to established analysis guidelines (Aquino et al., 2022; Parkes et al., 2018; Power, 2017). Probabilistic masks for GM, WM, and CSF estimated from anatomical data during fMRIPrep were spatially aligned to the BOLD image (Avants et al., 2008). To minimize partial volume effects (e.g., GM inclusion in the WM mask), the CSF mask was eroded once using a 3 x 3 x 3 mm kernel, whereas the WM mask was eroded up to five times. If an erosion iteration left fewer than 5 voxels in the mask, the previous erosion iteration was used. Mean time-series data within each of these eroded masks were then extracted from the preprocessed data (i.e., after any ICA-denoising, if applied). The first derivatives, their powers, and the powers of the raw time-series data were also calculated individually for the white matter, cerebrospinal fluid and gray matter tissue areas. These were then concatenated with the 24P motion parameters calculated in fMRIPrep to create participant-specific confound matrices. GMSR was performed with the associated expansion and derivative terms (four parameters total).

We also ran all of our pipelines with and without SR as a censoring strategy. Frames targeted for censoring were identified with respect to participant-specific framewise displacement (FD) measures (Yan et al., 2013). FD quantifies degree of head motion and is estimated from head-translation and rotation. To account for pseudomotion arising due to the rapid TR of the multiband sequence, we followed prior work (Fair et al., 2020) and filtered the six rotation and translation head motion parameters using a band-stop Butterworth filter [0.31–0.41 Hz]. Summed absolute successive differences were computed from these processed vectors and used to re-estimate FD for each time point (Fair et al., 2020). Timepoints exceeding 0.20mm in FD were flagged for spike regression. If fewer than 5 timepoints occurred between periods flagged for spike regression, these originally unmarked volumes were also included for spike regression (Gordon et al., 2014). 8P regression +/- GMSR (with and without SR) were performed in a single step and subsequent to any ICA-based denoising using the regression model implemented in fsl_regfilt (Lindquist et al., 2019; Satterthwaite et al., 2013).

### RIPTiDe

The toolbox Rapidtide was used to estimate low-frequency oscillations (LFO) via the Regressor Interpolation at Progressive Time Delays (RIPTiDe) algorithm (Tong et al., 2019) (https://github.com/bbfrederick/rapidtide). Time delay analyses were performed for each gray-matter voxel’s time series to assess similarity with a reference time series in the LFO band. More specifically, a “probe regressor” was derived from empirical data by filtering the average gray matter BOLD signal to frequencies within the LFO band (i.e., 0.01 Hz < *f* < 0.15 Hz) to capture hemodynamic fluctuations. Time-lagged voxelwise BOLD amplitude changes were then cross-correlated with this probe regressor to identify the most strongly correlated lag within a time window of +/-10 seconds, with probe-related (i.e. blood arrival related) variance then removed from the voxel time series using linear regression. This step was performed as an alternative to GMSR, prior to bandpass filtering. Pipelines involving 24P and ICA-AROMA can help improve LFO estimation by reducing motion artifacts without removing haemodynamic noise, but RIPTiDe is not validated for data denoised with 8P, ICA-FIX, or ME-ICA, which aim to mitigate sources of noise that may overlap with those targeted by RIPTiDe. In pipelines involving 8P, ICA-FIX, or ME-ICA, RIPTiDe regressors were estimated from the data with only 24P applied. The RIPTiDe regressors were then removed from the data in a separate stage to which 8P, ICA-FIX, or ME-ICA regressors were applied, as is the default.

### FC Estimation

The gray matter probability map derived from fMRIPrep and the preprocessed BOLD images were registered to MNI152NLin2009cAsym space (Avants et al., 2008) and multiplied together to limit partial volume effects and assign higher weight to grey matter voxels when extracting regional mean time series (see below). The resulting functional data were bandpass filtered (0.008 < 𝑓 < .08 Hz) before parcellation using the combined Schaefer400 and Melbourne Scale 2 Subcortical Atlas, dividing the cortex into 400 and the subcortex into 32 functionally homogenous regions (Schaefer et al., 2018; Tian et al., 2020).

The mean time-series for each region was extracted and individual-specific FC matrices were calculated using product-moment correlations between regional time courses, as implemented in Nilearn, based on Nipy (Brett et al., 2020; Gorgolewski et al., 2011). These correlation values were then transformed into *z*-scores using Fisher’s *r*-to-*z* transformation (Fisher, 1915).

### Quality Control Metrics (QCMs)

Six data quality evaluation metrics were assessed for each pipeline: (1) variance explained by the first principal component of the parcellated time-series (VE1); (2) delta variation signal (DVARS); (3) temporal signal-to-noise ratio (TSNR); (4) quality-control functional-connectivity (QC-FC); (5) QC-FC-Distance Dependence; and (6) functional connectivity inflation (FCI).

VE1 characterizes the extent to which the FC estimates are globally correlated. Higher values suggest strong WSD contamination, potentially arising from major fluctuations in respiration and/or head motion (Aquino et al., 2020). DVARS measures the similarity of voxel signal amplitude between consecutive time points (Afyouni & Nichols, 2018; Power et al., 2012). Averaging DVARS across the brain mask provides an overall metric of voxel signal homogeneity over time. High values indicate strong fluctuations in signal intensity over time, which may be induced by head motion. TSNR is estimated as the mean signal of a voxel over time divided by its standard deviation. Unlike DVARS, higher average TSNR values across the brain indicate greater stability, while lower values suggest more variability which may be driven by noise (Murphy et al., 2006).

QC-FC is estimated at each edge and quantifies the degree to which a given FC estimate correlates with a summary measure of head motion, quantified using mean FD, across participants (Aquino et al., 2020; Ciric et al., 2017; Parkes et al., 2018). The distribution of QC-FC values should have a mode of zero and minimal variance following the successful mitigation of motion-related artifacts. QC-FC Distance Dependence reflects the degree to which QC-FC relationships correlate with the Euclidean distances between each pair of regions. High head motion can drive higher FC estimates for short-range connections and lower FC for long-range connections (Ciric et al., 2017; Power et al., 2012).

Finally, FCI refers to an artefactual increase of FC as a function of time spent in the scanner (Korponay et al., 2024). Many contemporary preprocessing strategies are incapable of effectively removing this artefact (Korponay et al., 2023). Twenty consecutive windows (approximately 34 seconds each) were investigated for the present analyses. Previous research on FCI has used time-windows as short as 21.6s (Korponay et al., 2023), but larger window sizes (approaching 40 seconds) may allow for greater stability in estimates of dynamic changes in FC (Zalesky & Breakspear, 2015; Zhuang et al., 2020).

### Participant Exclusion

Sixty participants were removed on the basis functional data preprocessing failures, session interruption or incomplete fMRI data (i.e. fewer than 767 volumes collected), mean FD exceeding 0.25mm, >20% of FDs exceeding 0.20mm, any FDs exceeding 5.00mm, or if >50% of their data were flagged for spike regression (Aquino et al., 2020; Parkes et al., 2018; Pavlovich et al., 2024) (final *n* = 358).

#### Behavioral measures and predictions

The 60-item Big 5 Inventory II (BFI-II) measures the Big Five Personality Traits: Neuroticism, Extraversion, Conscientiousness, Agreeableness, and Openness. Participants respond on 5-point Likert scales from 1 (disagree strongly) to 5 (agree strongly) (Soto & John, 2017). Intelligence was also measured via the Wechsler Abbreviated Scale of Intelligence - Second Edition (WASI-II) (Wechsler, 1999, 2018). More specifically, we measured Vocabulary and Matrix Reasoning scores.

After exclusions, 4.64% of survey data were missing across the 358 retained participants and variables. Missing data in behavioral measures were imputed using predictive mean matching, a widely used statistical method generally associated with greater accuracy in missing data estimation relative to other imputation methods (Austin et al., 2021; Bailey et al., 2020; Morris et al., 2014). Imputation was conducted using the Multivariate Imputation by Chained Equations (MICE) approach implemented in the *mice* package (van Buuren & Groothuis-Oudshoorn, 2011) written for R (R Core Team, 2013). Although this approach typically generates multiple datasets with different imputations for missing values, a single imputed dataset was selected at random for input into subsequent KRR analyses to provide an almost unbiased estimate of the missing data.

## Results

### Evaluation of Pipeline-Level QCMs

We first evaluated six QCMs per pipeline: VE1, DVARS, TSNR, QC-FC, QC-FC Distance, and FCI. As expected, the removal of WSDs via either GMSR or RIPTiDE led to a dramatic reduction of VE1, with no obvious differences between SE and ME data or between different ICA denoising strategies (Figure 1). For DVARS and TSNR, ME pipelines showed a major advantage over SE pipelines. Applying ME-ICA with GMSR or RIPTiDE led to further improvements on these benchmarks. There was a marginal gain offered by combining ICA-FIX with ME-ICA compared to combining ME-ICA with ICA-AROMA (Figures 2 and 3). For SE data, ICA-based pipelines led to better DVARS and TSNR estimates compared to non-ICA-based pipelines, with ICA-FIX performing better than ICA-AROMA.

**Figure 1.**
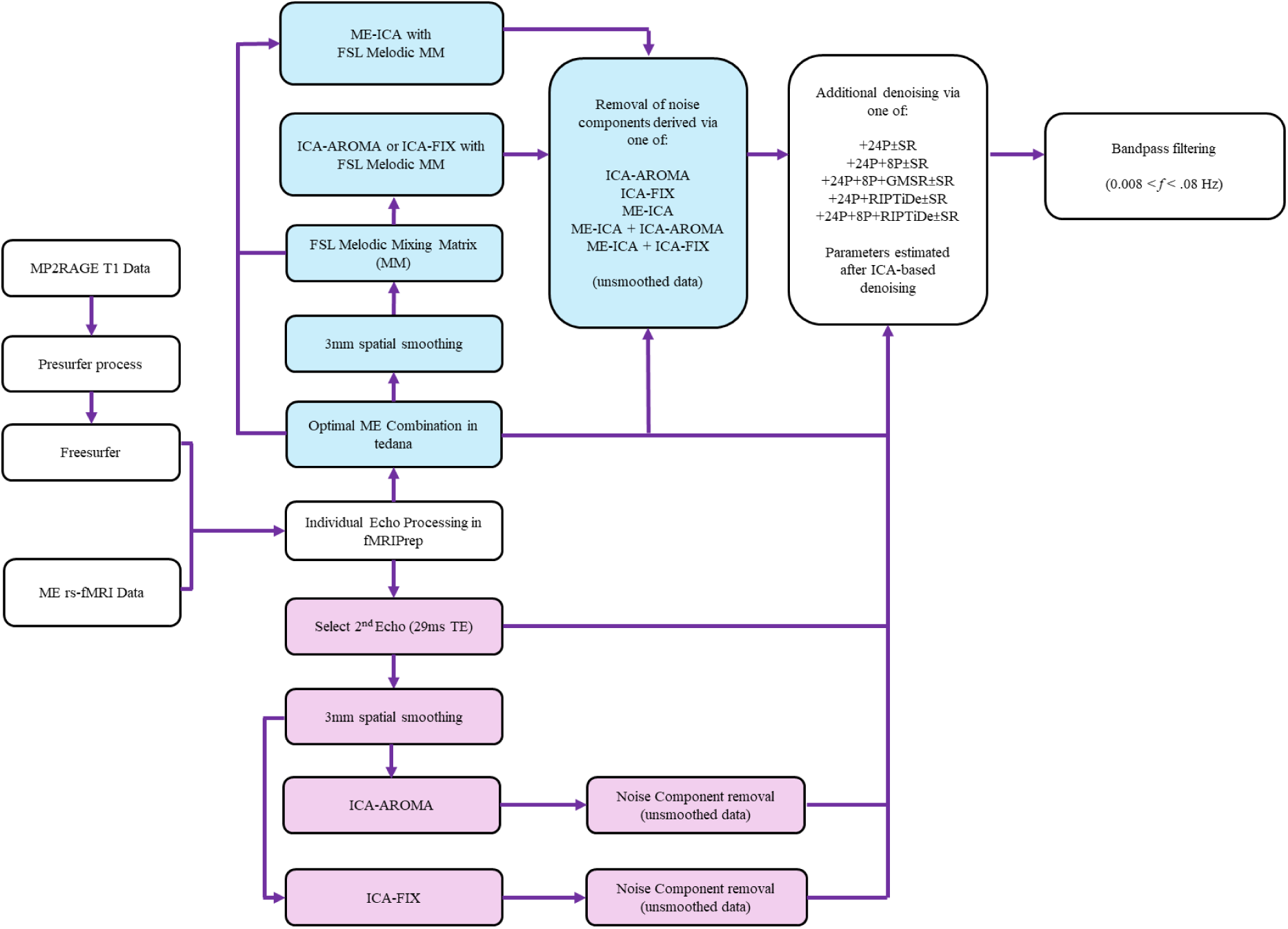
Overview of data preprocessing pipelines. Multi-echo-specific preprocessing steps are coloured light blue; single-echo-specific preprocessing steps are coloured pink. Common preprocessing steps are in white boxes. ME – multi-echo; FSL - FMRIB Software Library; ME-ICA – Multi-Echo-Independent Components Analysis; ICA-FIX - Independent Component Analysis - FMRIB’s Independent Component Analysis-based X-noiseifier; ICA-AROMA – Independent Component Analysis Automatic Removal of Motion Artifacts; MM – mixing matrix; 24P - raw head rotation and translation information, their first-order derivatives, their quadratics, and the quadratics of their derivatives; 8P - average cerebrospinal fluid and white matter signals, their derivatives, their quadratics, and the quadratics of their derivatives; GMSR - average gray matter signals, their derivatives, their quadratics, and the quadratics of their derivatives; SR – spike regression – a strategy for volume censoring.

**Figure 2.**
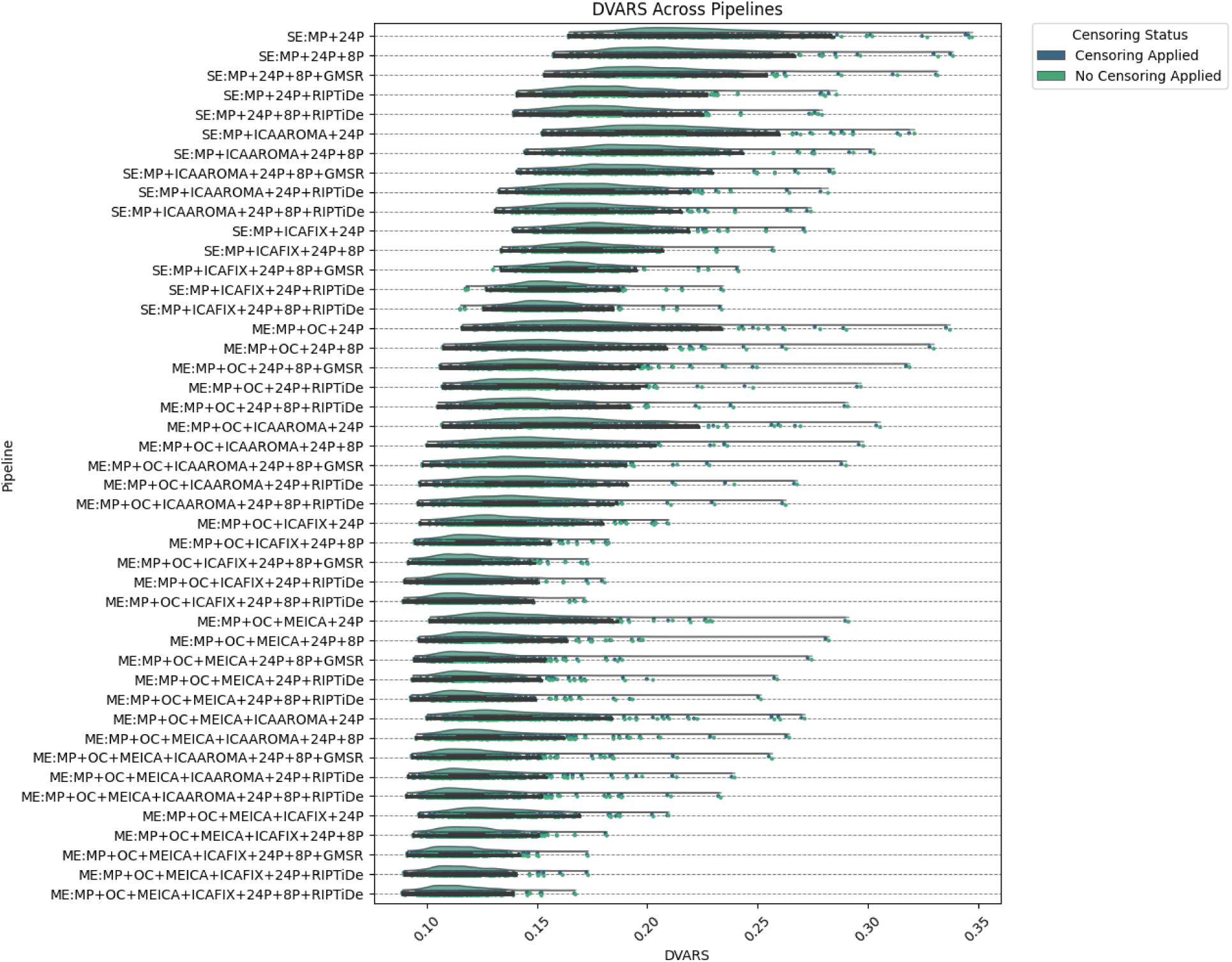
The effect of various denoising strategies on subject-level DVARS, aggregated at the pipeline level. Smaller values indicate less noise. Note that histograms for Censoring Status (applied or not applied) overlap substantially. *MP* refers to minimal preprocessing with fMRIPrep.

**Figure 3.**
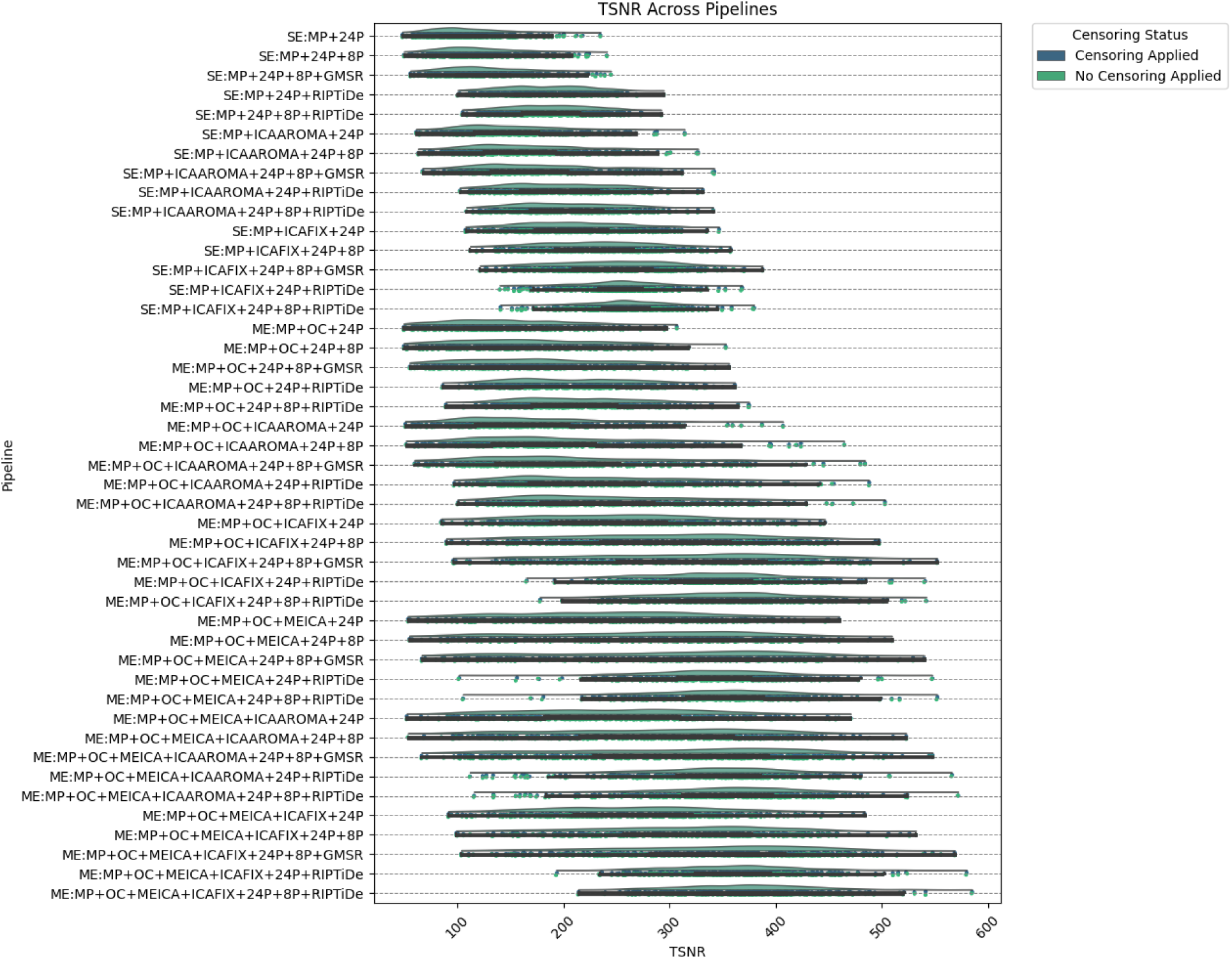
The effect of various denoising strategies on subject-level TSNR, aggregated at the pipeline level. Larger values indicate less noise. Note that histograms for Censoring Status (applied or not applied) overlap substantially. *MP* refers to minimal preprocessing with fMRIPrep.

Pipelines with some form of WSD removal (i.e., GMSR or RIPTiDE) improved QC-FC correlation (*ρ*) distributions (i.e., the mode of the distribution was closer to zero; Figure 4) and reduced the number of edges demonstrating significant relationships (𝑝 < .05, uncorrected) with head motion (Figure 5). We also observed a consistent, yet minor, gain for GMSR over RIPTiDE across SE and ME pipelines for these metrics. QC-FC distance dependence was more variable overall, ranging between 𝜌 = -.18 (for ME:MP+OC+24P+8P+GMSR) and 𝜌 = .07 (for ME:MP+OC+MEICA+ICAFIX+24P+RIPTiDe) (Figure 6). In SE data, the best-performing pipelines for reducing distance-dependence used ICA-FIX. In ME data, the application of ME-ICA generally improved distance-dependence relative to pipelines that did not use ME-ICA, with the best-performing ME pipelines combining ME-ICA, ICA-FIX and GMSR, or combining ME-ICA, ICA-AROMA, and GMSR. The application of RIPTiDE instead of GMSR in these pipelines led to a slight worsening of QC-FC distance-dependence (e.g., from 𝜌 = -.01 for pipeline ME:MP+OC+MEICA+ICAAROMA+24P+8P+RIPTiDe to 𝜌 = .07 for pipeline ME:MP+OC+MEICA+ICAFIX+24P+RIPTiDe). FCI was evident in both SE and ME data and was generally only mitigated when some form of WSD removal (GMSR or RIPTiDE) was used (Figure 7). Censoring was not associated with major variations in any of the data quality metrics.

**Figure 4.**
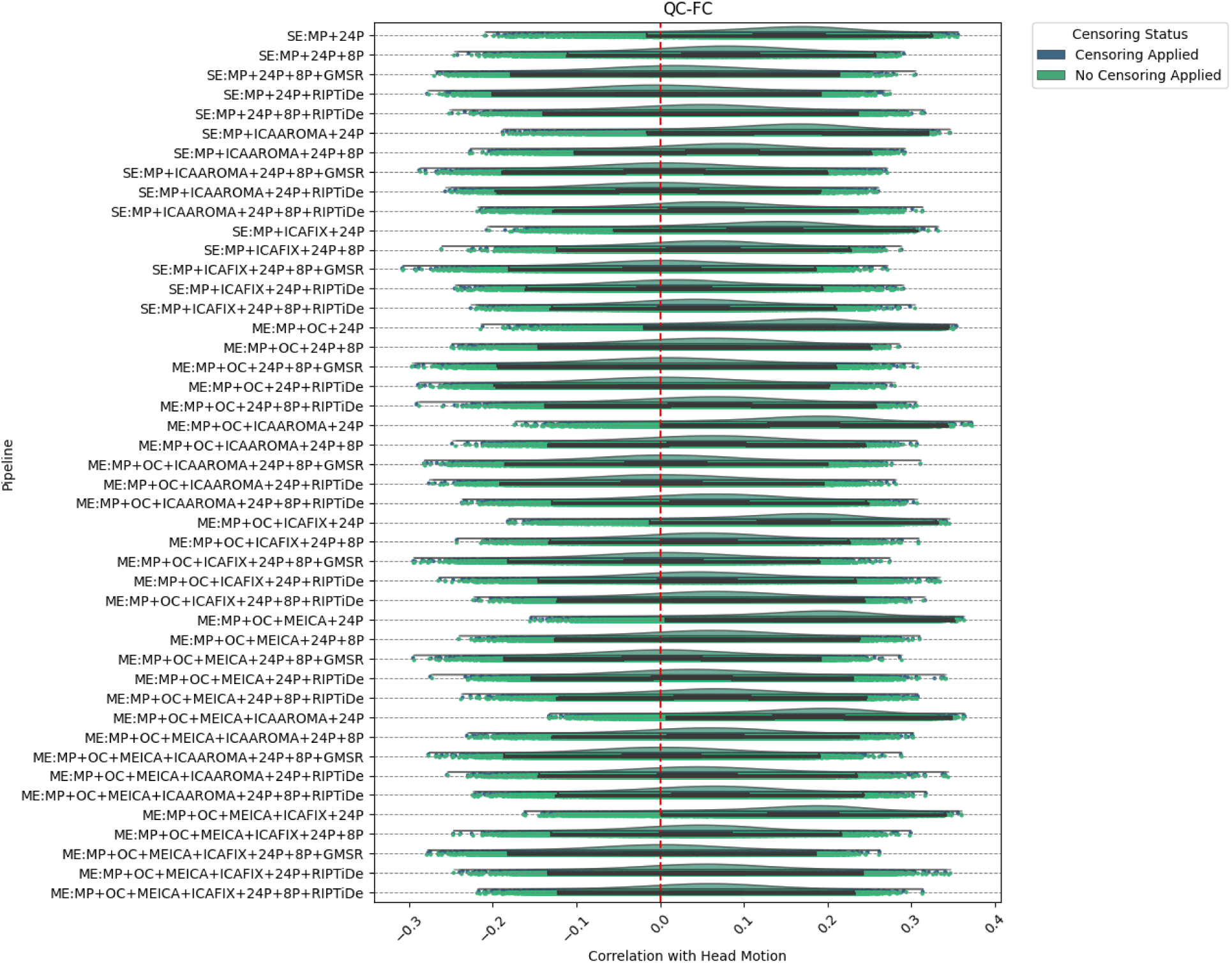
The effect of various denoising strategies on subject-level QC-FC, aggregated at the pipeline level. Correlations are Spearman’s rho. Distributions centered at zero indicate less head-motion contamination. Note that histograms for Censoring Status (applied or not applied) overlap substantially. *MP* refers to minimal preprocessing with fMRIPrep.

**Figure 5.**
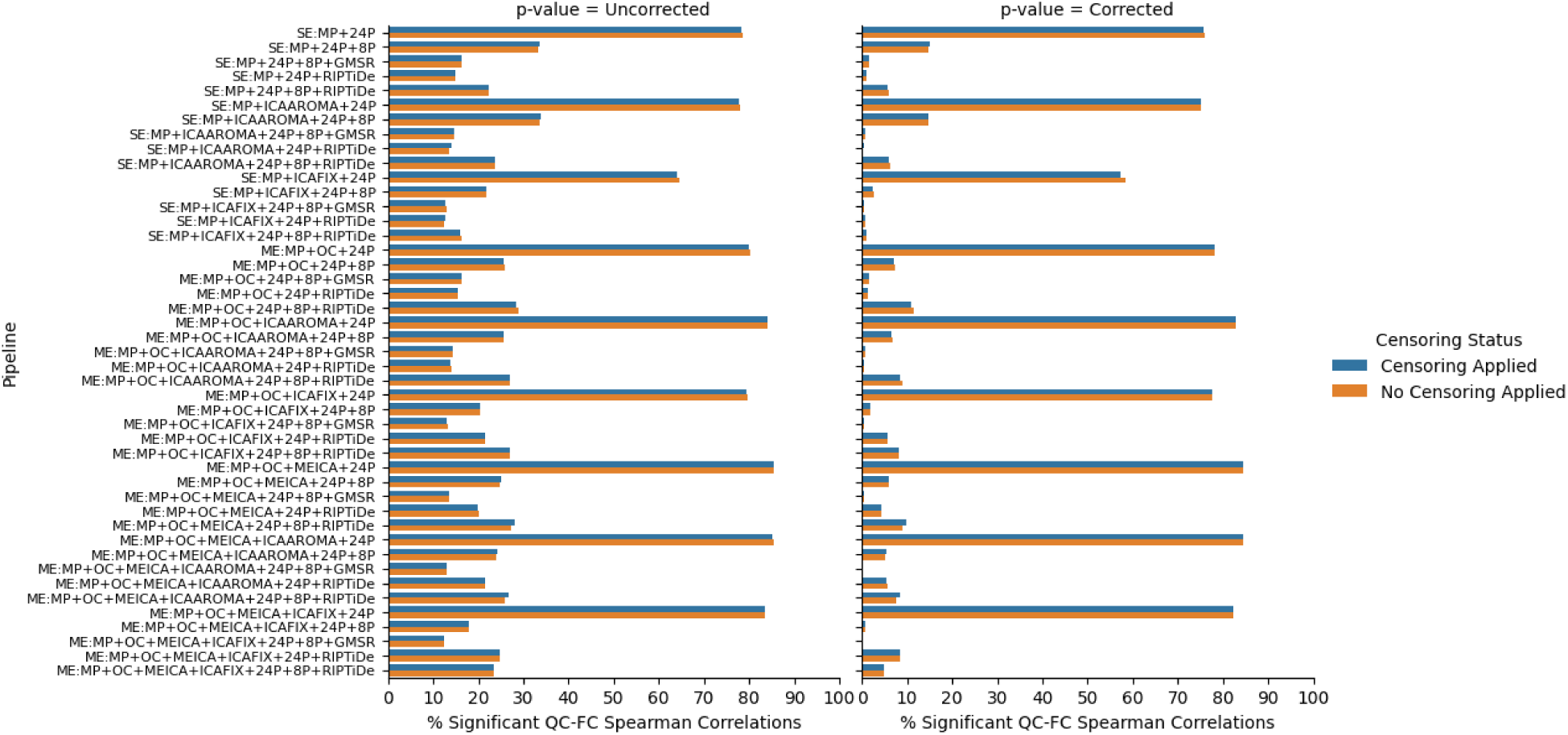
The effect of various denoising strategies on the number of statistically significant QC-FC values (uncorrected values displayed left; Benjamini-Hochberg corrected *p-*values right). A larger overall percentage of edges significantly correlated with head motion indicates greater amounts of head-motion contamination in the data. *MP* refers to minimal preprocessing with fMRIPrep.

**Figure 6.**
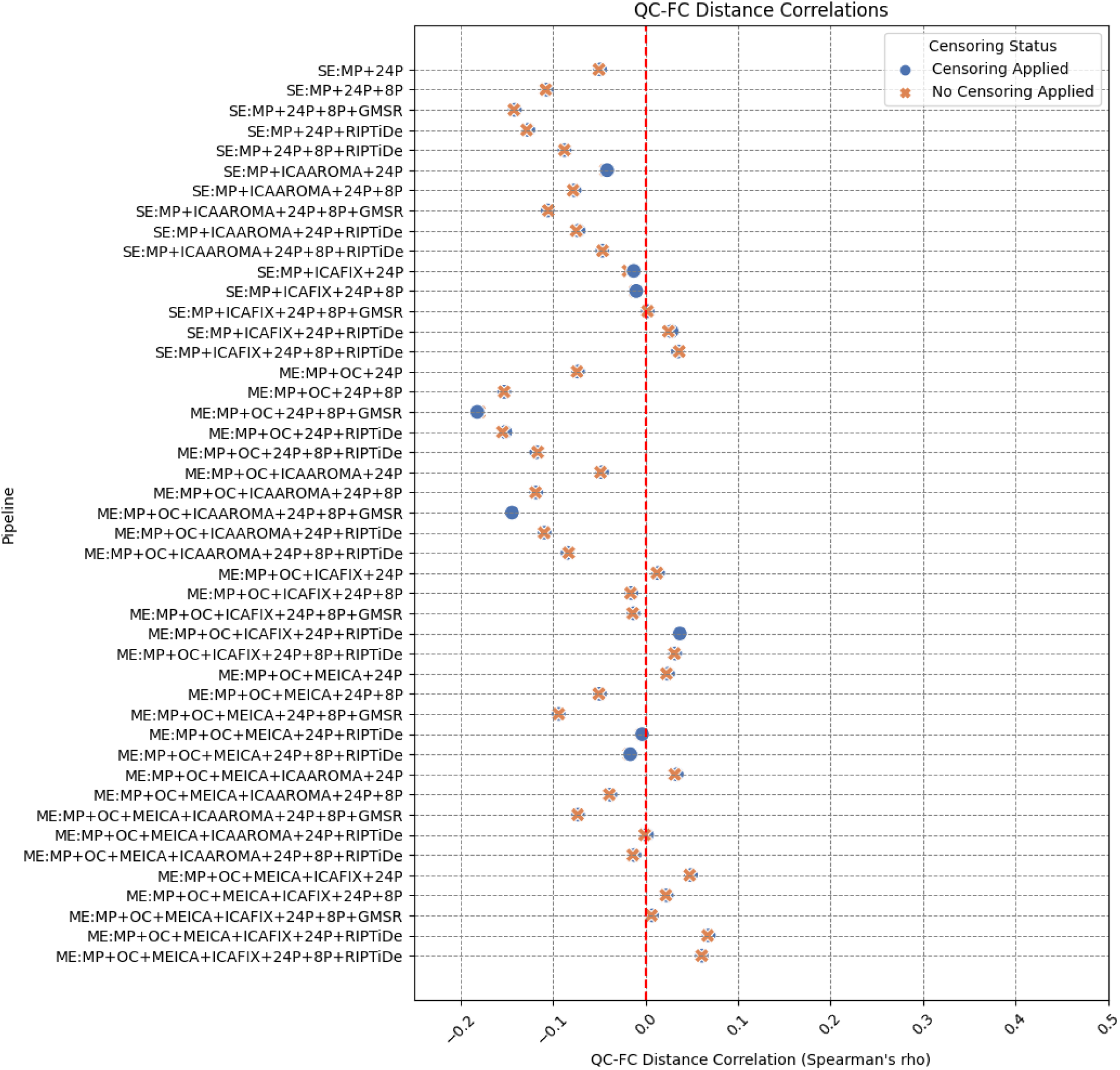
The effect of various denoising strategies on pipeline-level QC-FC Distance Dependence. Correlations are Spearman’s rho (ρ). Distributions centered at zero indicate less head-motion contamination in the data. Note that points for Censoring Status (applied or not applied) overlap substantially. *MP* refers to minimal preprocessing with fMRIPrep.

**Figure 7.**
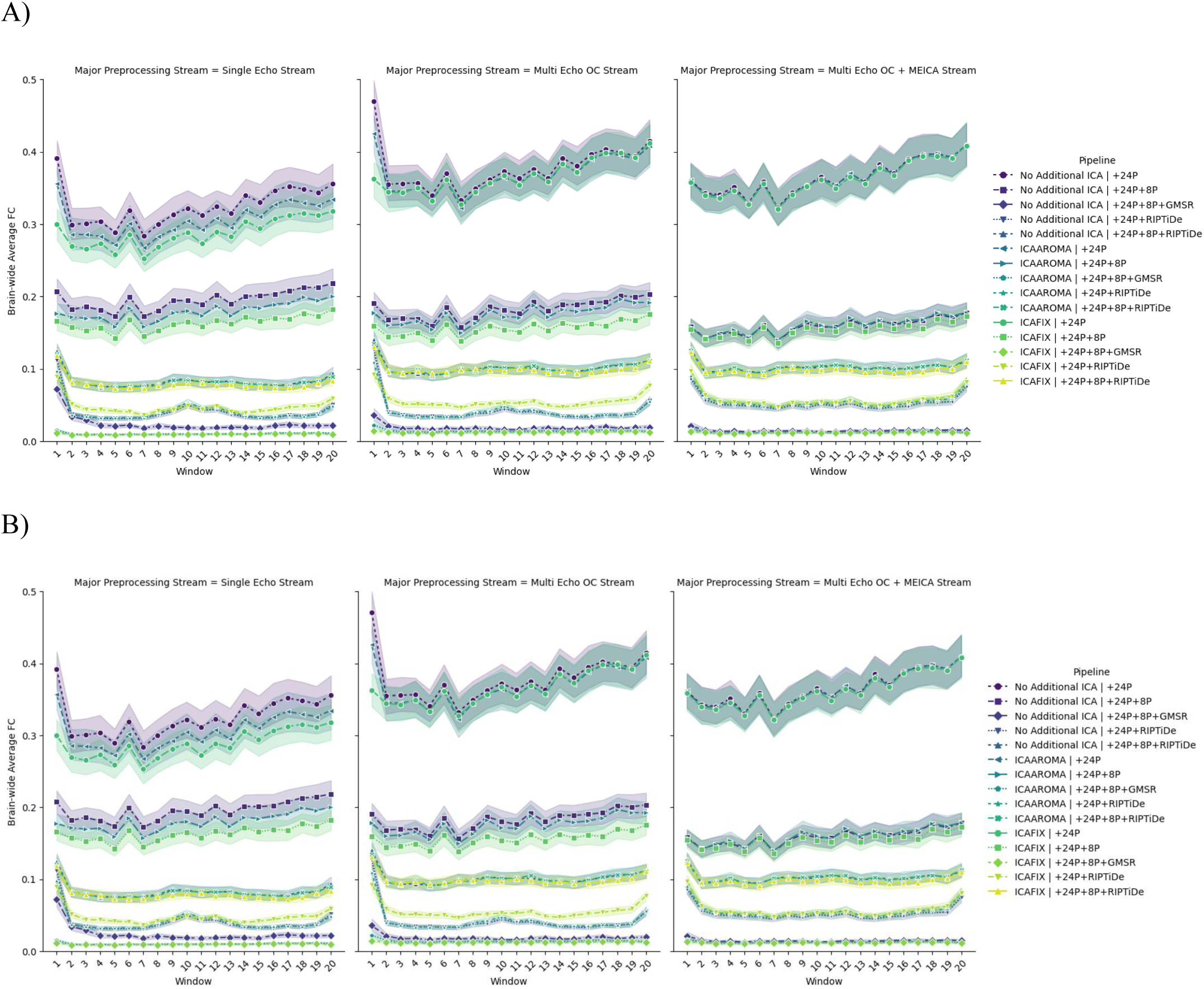
The effect of various denoising strategies on subject-wise FCI for single and multi-echo data, aggregated at the pipeline level. Censoring was applied for pipelines in Panel A. No censoring was applied for Panel B. Individual time series were discretized into twenty windows. Lack of line slope indicates diminished FCI. No trend was observed for Censoring Status, or whether SE data were used or whether OC or ME-ICA was applied. Note that Additional ICA Denoising categories overlap considerably.

### Behavioural predictions

We next evaluated behavioural prediction accuracies using cross-validated FC-based KRR (J. Li et al., 2019). Aggregating results across both cognitive and personality measures via averaging (Figure 10), the best performance was achieved for ME pipelines that did not use WSD removal. Specifically, four ME pipelines stood out as having the highest accuracies: ME:MP+OC+ICA-FIX+24P, ME:MP+OC+ME-ICA+24P, ME:MP+OC+ME-ICA+ICA-FIX+24P, ME:MP+OC+ME-ICA+ICA-AROMA+24P. Of ME pipelines including WSD removal, pipelines combining ICA-AROMA without ME-ICA showed higher average performance than those using ME-ICA or ICA-FIX. There was a slight performance disadvantage for pipelines combining ME-ICA, ICA-FIX, and WSD removal, suggesting that these pipelines may be overly aggressive and remove behaviorally relevant variance despite their advantages on QCMs.

Of the pipelines that were not ranked in the top 4, SE pipelines generally showed higher average performance than ME pipelines, with some evidence that WSD removal led to decreased predictive accuracy. Similar trends were observed when inspecting results for cognition and personality separately (Figures 8 and 9). Censoring had minimal effect on model accuracies. Note that while performance differences between some pipelines exceeded 100%, prediction accuracies were generally small and never exceeded 𝑟 = 0.15.

**Figure 8.**
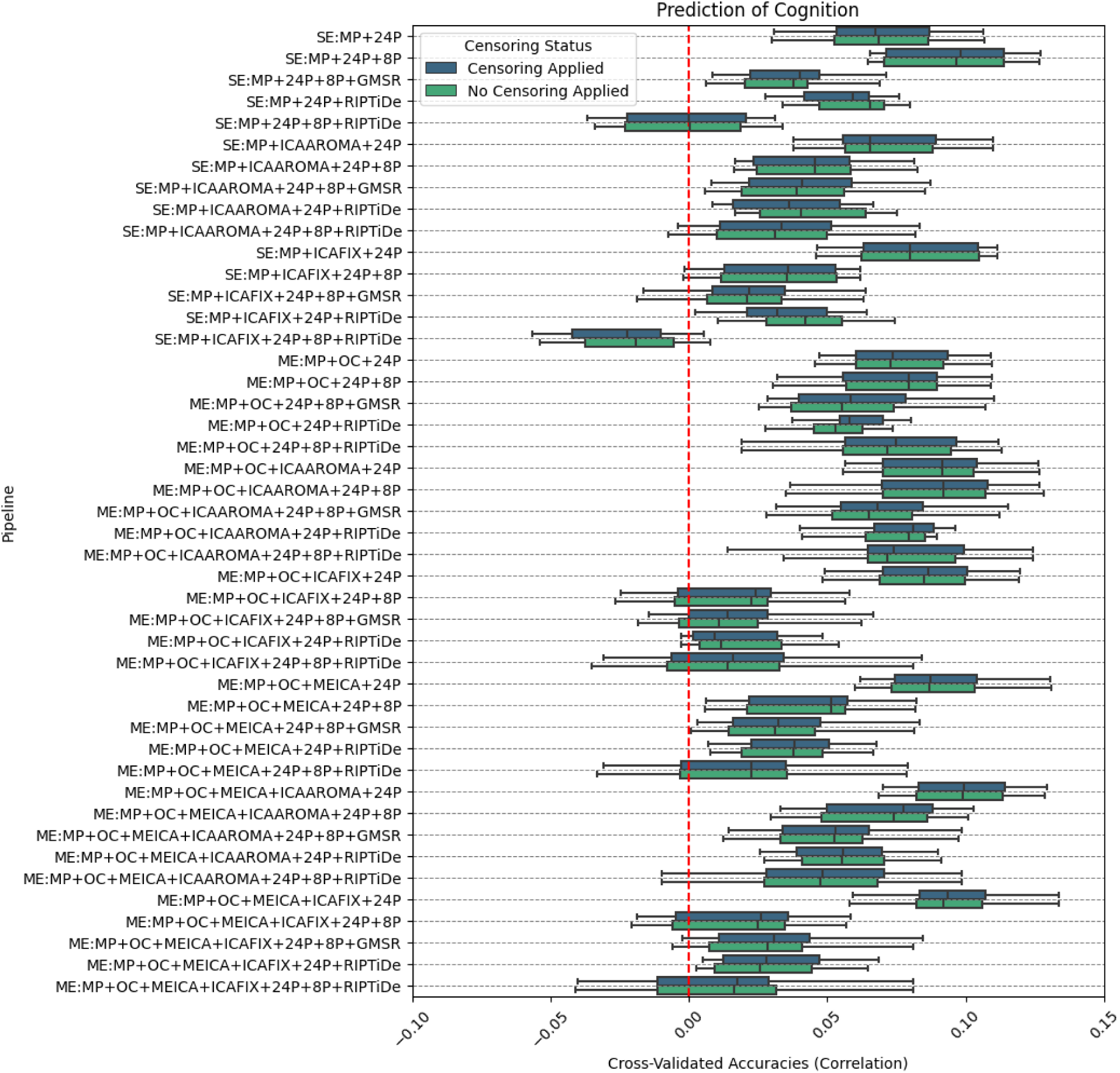
The effect of various denoising strategies on pipeline-level cognition prediction. Cognitive measures were individual-level *T*-score transformed WASI-II Matrix Reasoning and WASI-II Vocabulary scores, averaged across the two tasks. Pipelines are ordered based on the purported level of denoising aggressivity/complexity. The results reported here are typical of machine learning neuroimaging studies. *MP* refers to minimal preprocessing with fMRIPrep.

**Figure 9.**
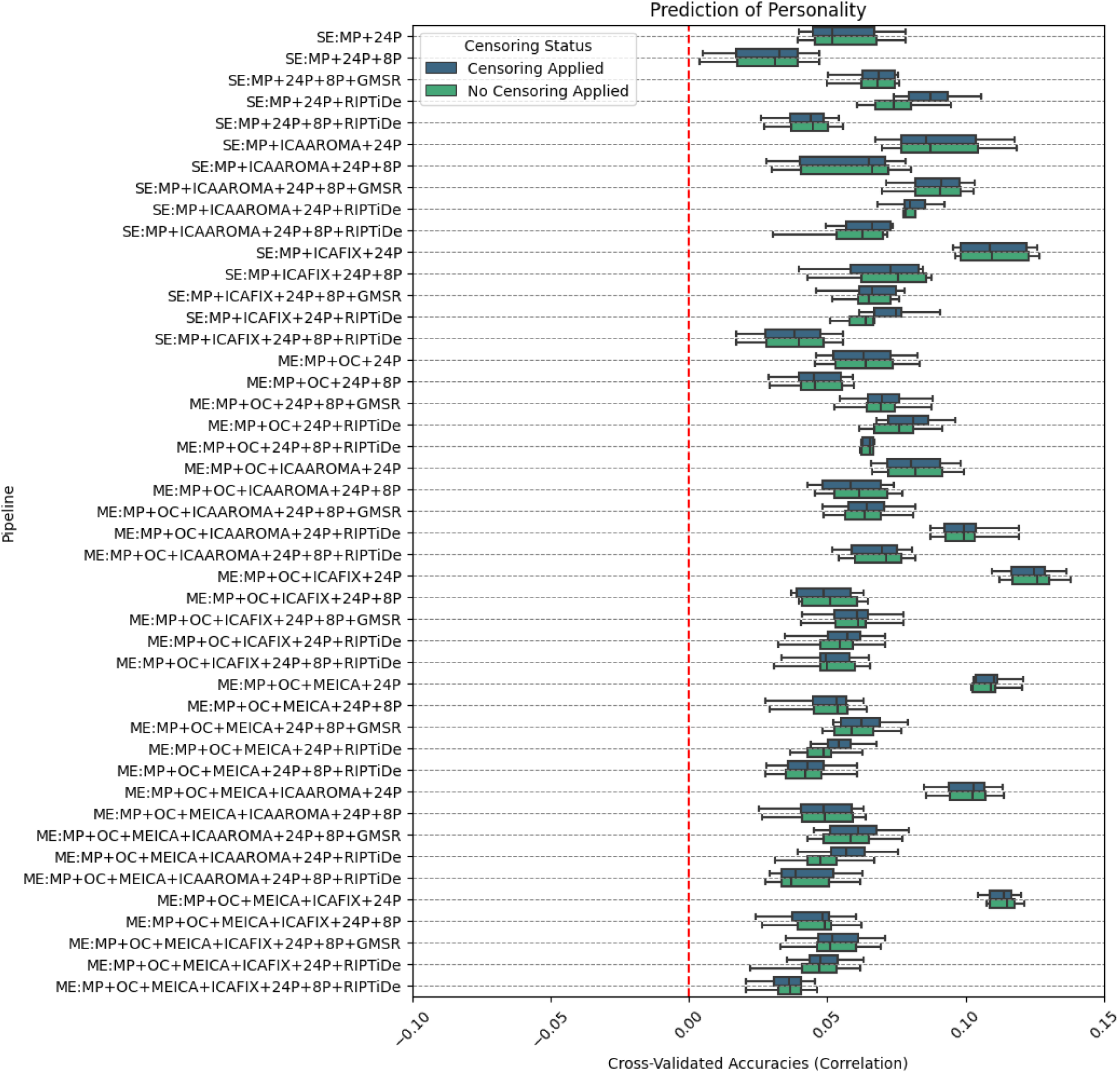
The effect of various denoising strategies on pipeline-level personality prediction. Personality measures were individual-level Big Five (domain) scores. Pipelines are ordered based on the purported level of denoising aggressivity/complexity. The results reported here are typical of machine learning neuroimaging studies. *MP* refers to minimal preprocessing with fMRIPrep.

**Figure 10.**
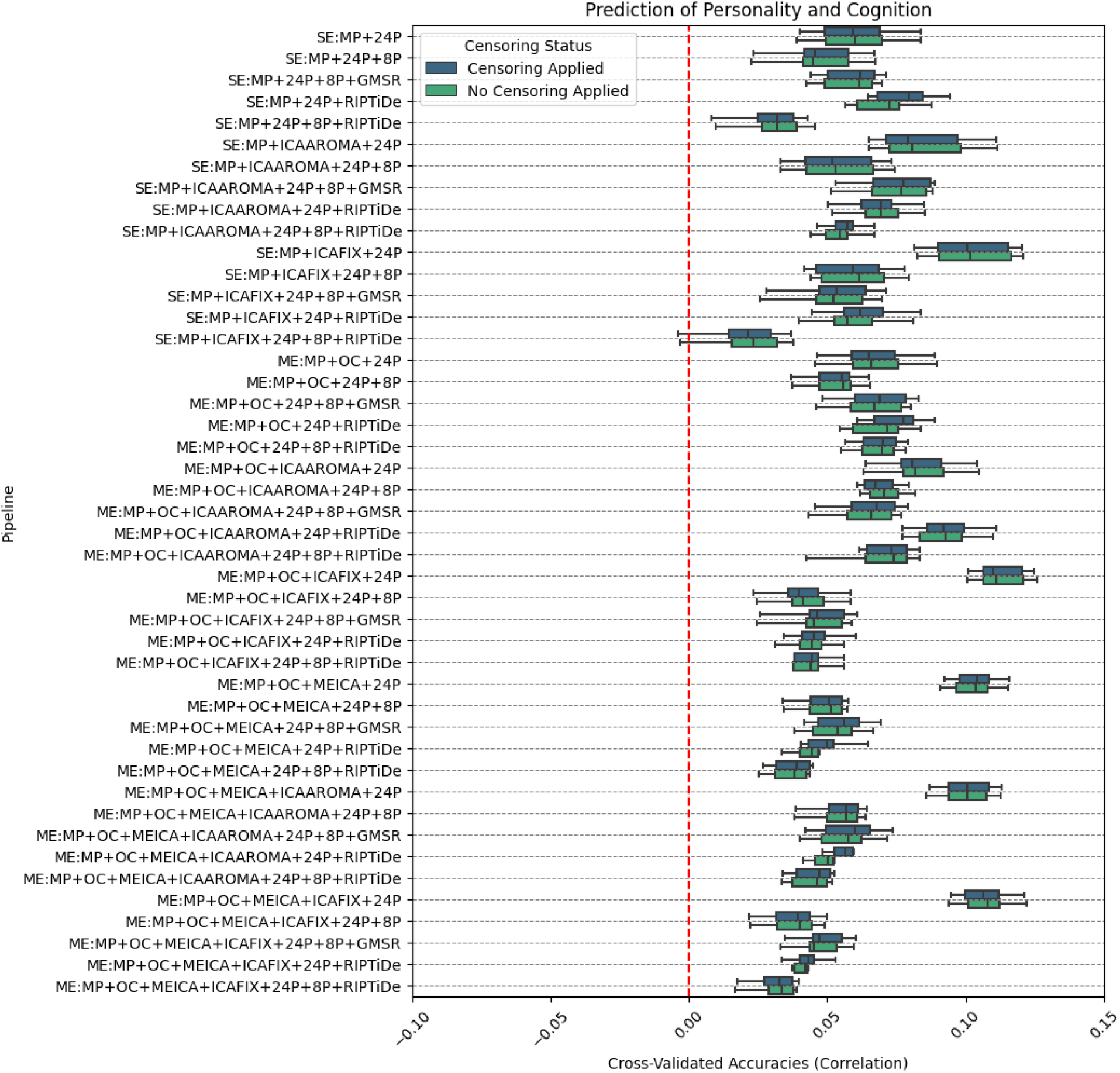
The effect of various denoising strategies on pipeline-level personality and cognition prediction. Personality measures were individual-level Big Five (broad domain) scores, and cognitive measures were individual-level *T*-score transformed WASI-II Matrix Reasoning and WASI-II Vocabulary scores. Pipelines are ordered based on the purported level of denoising aggressivity/complexity. The results reported here are typical of machine learning neuroimaging studies. *MP* refers to minimal preprocessing with fMRIPrep.

### Pipeline Rankings

We ranked each pipeline based on its performance across multiple metrics using two methods: ordinal rankings and percentage decrement. In the ordinal ranking scheme, we assigned a rank of 1 to the top-performing pipeline and progressively ranked other pipelines in order with higher numbers. As such, lower values indicated better performance. Since it is difficult to quantify and rank pipelines in their degree of FCI improvement, we simply assigned a score of 1 to pipelines that removed FCI (i.e., pipelines using RIPTiDe or GMSR) and a score of 2 to all others. In the percentage decrement ranking scheme, the best performing pipeline was assigned a score of 100% and all other pipelines were ranked as a percentage decrement relative to this value (with lower percentages indicating worse performance). For FCI, pipelines incorporating the RIPTiDe algorithm or GMSR were assigned a score of 100% and all others received a score of 0%. To obtain an overall rank for each pipeline’s denoising efficacy, we averaged the rankings across denoising metrics from both the ordinal and percentage decrement schemes independently. The same approach was applied to KRR results (i.e., averaging ranks across Figures 8 and 9 independently before combination), yielding overall ranks for behavioural predictions. These ranks were then averaged to obtain an overall performance ranking for each pipeline. Note that these rankings weight QC and behavioural performance benchmark metrics equally and should thus be considered as heuristics, as different investigators may wish to prioritize specific benchmarks in certain studies. The top non-censored ME and SE rankings (according to ordinal rankings) are shown in Table 1; full results can be found in Supplementary Material 1. The rankings were largely the same when censoring was incorporated into the pipeline. The ME:MP+OC+ICAAROMA+24P+RIPTiDe was the best pipeline according to both ranking schemes. Excluding the binary ranking associated with FCI removal did not change the finding.

**Table 1.** Top 5 (non-censored) ME and SE pipelines with regards to the Overall Performance composite, displayed in ascending order (according to ordinal rankings). This was computed via the average of Overall Denoising Efficacy and Overall Behavioural Prediction rank. Censoring did not appear to make a difference with regards to pipeline performance at this coarse level of measurement. *MP* refers to minimal preprocessing with fMRIPrep.

|  | Pipeline | Overall Ordinal Rank | Overall Percentage Decrement |
| --- | --- | --- | --- |
| *Top 10 ME Pipelines* | ME:MP+OC+ICAAROMA+24P+RIPTiDe | 24.39 | 75.90% |
|  | ME:MP+OC+ICAFIX+24P | 27.89 | 63.49% |
|  | ME:MP+OC+MEICA+ICAFIX+24P | 28.54 | 61.20% |
|  | ME:MP+OC+MEICA+ICAAROMA+24P | 29.29 | 59.18% |
|  | ME:MP+OC+MEICA+24P | 30.64 | 58.84% |
|  | ME:MP+OC+MEICA+ICAAROMA+24P+8P+GMSR | 34.50 | 64.22% |
|  | ME:MP+OC+ICAAROMA+24P+8P+RIPTiDe | 34.93 | 67.22% |
|  | ME:MP+OC+24P+RIPTiDe | 35.96 | 61.56% |
|  | ME:MP+OC+ICAAROMA+24P+8P+GMSR | 37.25 | 62.09% |
|  | ME:MP+OC+MEICA+ICAFIX+24P+8P+GMSR | 37.68 | 61.16% |
| *Top 10 SE Pipelines* | SE:MP+ICAFIX+24P | 32.25 | 59.52% |
|  | SE:MP+ICAFIX+24P+RIPTiDe | 33.04 | 62.47% |
|  | SE:MP+ICAAROMA+24P+RIPTiDe | 33.82 | 62.26% |
|  | SE:MP+24P+RIPTiDe | 35.25 | 62.15% |
|  | SE:MP+ICAAROMA+24P+8P+GMSR | 36.93 | 60.90% |
|  | SE:MP+ICAFIX+24P+8P+GMSR | 37.39 | 58.06% |
|  | SE:MP+ICAAROMA+24P | 41.04 | 44.53% |
|  | SE:MP+ICAFIX+24P+8P | 41.75 | 50.74% |
|  | SE:MP+24P+8P+GMSR | 44.86 | 50.39% |
|  | SE:MP+ICAAROMA+24P+8P+RIPTiDe | 47.96 | 54.17% |

## Discussion

The optimum approach for denoising fMRI data remains an open area of inquiry. In this study, we compared 90 different SE and ME pipelines with respect to 6 different measures of data quality in addition to evaluating their influence on the prediction of behavioral measures measured outside the scanner. We determined that ME pipelines generally outperformed SE pipelines across our denoising metrics and also yielded the highest effect sizes for behavioural prediction models. An approximate ranking scheme aggregating results across all benchmarks indicated that ME pipelines involving ICA-based denoising generally fared well, with ME:MP+OC+ICAAROMA+24P+RIPTiDe offering the best trade-off between denoising efficacy and behavioural prediction.

### Benchmarking denoising efficacy

The use of ME data was associated with performance advantages for DVARS, TSNR, and QC-FC distance-dependence, consistent with past findings (Kundu et al., 2017). ME pipelines were associated with a slight increase in VE1 compared to their SE counterparts with no discernible differences in QC-FC correlations. These findings suggest that while ME acquisitions can improve SNR, they do not necessarily lead to a major improvement in the degree to which FC estimates are contaminated by motion.

For SE data, ICA-FIX outperformed ICA-AROMA and pipelines that did not use any ICA-based denoising strategy across several data quality metrics (TSNR, DVARS, QC-FC Distance Dependence), consistent with previous evidence that ICA-AROMA is a suboptimal denoising algorithm relative to ICA-FIX (Carone et al., 2017; Glasser et al., 2019). Among pipelines using ICA-FIX, the further application of RIPTiDe without additional white matter and cerebrospinal fluid regressors was generally associated with the best performance for VE1, TSNR, and DVARS, marginally surpassing the performance of GMSR. GMSR showed a small advantage over RIPTiDe for QC-FC. Only RIPTiDe and GMSR were successful in mitigating FCI.

In ME data, we also observed an advantage for pipelines using ICA-FIX over ME-ICA alone, ICA-AROMA and no s-ICA denoising for TSNR, DVARS, and QC-FC Distance Dependence. However, the best denoising performance resulted from the combination of ME-ICA and ICA-FIX. The additional use of either GMSR or RIPTiDE led to further performance gains, with the differences between the two forms of WSD removal being marginal.

Together, these findings suggest that ME pipelines improve SNR and that the combination of ME-ICA, ICA-FIX, and either RIPTiDE or GMSR results in the highest denoising efficacy. For SE data, ICA-FIX combined with either GMSR or RIPTiDE results in the best performance.

### Benchmarking behavioural prediction accuracy

QCMs do not quantify the degree of behaviourally relevant variance removed during denoising. As such, a pipeline may perform well on QCM-related benchmarks simply because it is too aggressive, removing both variance relevant for specific research questions and noise. We therefore also investigated relative performance in predicting cognitive and personality measures using KRR. These analyses suggested that the more aggressive pipelines may indeed remove behaviourally relevant signal. Specifically, while the best performing pipelines according to QCMs generally incorporated GMSR or RIPTiDE, the pipelines that were most predictive of behaviour did not involve WSD removal.

Previous work has demonstrated that head-motion covaries with interindividual differences in behavior, and that the removal of motion-associated variance from FC can reduce brain-behaviour relationships (Siegel et al., 2017), while other work has shown that WSD attenuation through GSR, which covaries with head motion, can boost brain-behavior relationships (J. Li et al., 2019). These results are further complicated by evidence that GSR may only enhance brain-behavior estimates in certain datasets (Pavlovich et al., 2024). Our analyses covaried for mean FD, suggesting that the relative success of pipelines that do not incorporate WSD removal is unlikely to be explained by residual covariance between head motion and behaviour. Together, these mixed findings suggest that behavioural prediction accuracy on its own may be an insufficiently sensitive measure of the ability of a pipeline to preserve useful signal and more sophisticated methods may be required (e.g., Aquino et al., 2020; Glasser et al., 2018).

Recent work indicates that FC estimates derived from task-based fMRI-derived FC more accurately predict behavior than resting-state FC estimates (Zhao et al., 2023). As such, resting-state designs may not provide optimal conditions for comparing predictive models of individual differences in behaviour. Although relatively under-acknowledged, measurement error in estimating interindividual differences in behaviour also imposes a significant limit on achieving robust and reliable brain-behavior associations. The adoption of statistical and psychometric approaches for deriving more precise estimates of behavior may help improve effect sizes (Tiego et al., 2023). Indeed, the prediction accuracies we observed were generally small (i.e., all *r*<.15) and differences between the top 4 pipelines and the next best-performing pipelines were on the order of *r*<.05, so the gains associated with choosing one particular pipeline over another are modest.

### A trade-off between noise removal and signal preservation?

No single pipeline simultaneously maximised data quality and behavioural prediction accuracy, which aligns with past work in SE datasets (Pavlovich et al., 2024). To discern overall pipeline performance, we generated overall rankings for the quality control metrics and behavioural prediction independently, before subsequently computing an overall ranking from these composites (see Table 1). We determined that SE pipelines generally underperformed relative to ME pipelines on these measures. We propose that ME sequences may present a more effective approach than SE sequences for studying brain-behavior associations with rs-fMRI while balancing denoising concerns. Optimal combination of ME data followed by ICA-AROMA, 24P regression and RIPTiDe, generally provided strong denoising efficacy while maintaining robust brain-behavior associations for ME data. However, we emphasize that our ranking scheme should be used as a heuristic only, and that investigators should consider whether they prioritize optimizing certain benchmarks over others.

### Limitations

Further work is needed to validate our tentative recommendations using independent datasets, particularly in light of research highlighting variability in pipeline performance across different datasets (Pavlovich et al., 2024). Future studies should investigate the factors contributing to variability in prediction and denoising performance across pipelines in different datasets.

Our study used a common minimal pre-processing pipeline. Recent evidence indicates that variations in these earlier steps can also impact FC estimates (Li et al., 2024). Understanding how choices at early processing stages interatc with denoising strategies will be an important extension of this work.

We used common mixing matrices derived from FSL for all ICA approaches, which allow used to easily combine different ICA-based approaches (e.g, ME-ICA and ICA-FIX). This implementation is distinct from the default approach in *tedana*, and should thus be considered when interpreting our findings. A further consideration is that ICA-based approaches incorporate a stochastic element that can yield variable results from run to run (Pavlovich et al., 2024). We did not investigate this variability here to limit computational burden, but it should be considered in any future applications.

## Conclusions

We compared denoising approaches for multi-echo (ME) and single-echo (SE) fMRI acquisitions, along with various techniques to manage widespread signal deflections (e.g., GMSR, RIPTiDe) following spatial ICA. We tested 90 different pipelines across six established data quality metrics and evaluated their capacity to predict personality and cognition using resting state functional connectivity (FC). Our results indicated that no single pipeline maximizes efficacy and behavioural prediction accuracies. However, balancing both quality control metric results and relative behavioral prediction performance, the application of ICA-AROMA, 24P regression, and RIPTiDe to ME data resulted in the best aggregate outcomes. In general, ME acquisitions appear superior to SE data.

## Author contributions

TC, JT, and AF conceptualised the study, interpreted the results, and wrote the manuscript. TC prepared and analysed the data. KP, AS, and PTL reviewed the code for data analysis. JT, NOT, BH, RO, JK, KF, KC, SB, JM, MB, and AF contributed to data acquisition. AF was supported by the Australian Research Council (ID: FL220100184) and National Health and Medical Research Council (ID: 1197431).

## Code availability

Project code can be found here: https://github.com/cicadawing/Single-vs-Multi-Echo-fMRI-Denoising-Strategies

